# A machine learning approach to optimizing cell-free DNA sequencing panels: with an application to prostate cancer

**DOI:** 10.1101/2020.04.30.069658

**Authors:** Clinton L. Cario, Emmalyn Chen, Lancelote Leong, Nima C. Emami, Karen Lopez, Imelda Tenggara, Jeffry P. Simko, Terence W. Friedlander, Patricia S. Li, Pamela L. Paris, Peter R. Carroll, John S. Witte

## Abstract

**Background:** Cell-free DNA’s (cfDNA) use as a biomarker in cancer is challenging due to genetic heterogeneity of malignancies and rarity of tumor-derived molecules. Here we describe and demonstrate a novel machine-learning guided panel design strategy for improving the detection of tumor variants in cfDNA. Using this approach, we first generated a model to classify and score candidate variants for inclusion on a prostate cancer targeted sequencing panel. We then used this panel to screen tumor variants from prostate cancer patients with localized disease in both *in silico* and hybrid capture settings.

**Methods:** Whole Genome Sequence (WGS) data from 550 prostate tumors was analyzed to build a targeted sequencing panel of single point and small (<200bp) indel mutations, which was subsequently screened *in silico* against prostate tumor sequences from 5 patients to assess performance against commonly used alternative panel designs. The panel’s ability to detect tumor-derived cfDNA variants was then assessed using prospectively collected cfDNA and tumor foci from a test set 18 prostate cancer patients with localized disease undergoing radical proctectomy.

**Results:** The panel generated from this approach identified as top candidates mutations in known driver genes (e.g. HRAS) and prostate cancer related transcription factor binding sites (e.g. MYC, AR). It outperformed two commonly used designs in detecting somatic mutations found in the cfDNA of 5 prostate cancer patients when analyzed in an *in silico* setting. Additionally, hybrid capture and 2,500X sequencing of cfDNA molecules using the panel resulted in detection of tumor variants in all 18 patients of a test set, where 15 of the 18 patients had detected variants found in multiple foci.

**Conclusion:** Machine learning-prioritized targeted sequencing panels may prove useful for broad and sensitive variant detection in the cfDNA of heterogeneous diseases. This strategy has implications for disease detection and monitoring when applied to the cfDNA isolated from prostate cancer patients.

## Background

Substantial research has explored potential oncological applications of cell-free DNA (cfDNA), including in early detection, monitoring of residual disease, recurrence following treatment, and as a discovery tool for determining actionable therapeutic targets [1–3]. However, success using cfDNA in cancer has been limited by heterogeneity and signal intensity. In the context of heterogeneous cancers like those of the prostate, cfDNA also provides an opportunity to comprehensively measure tumor clonality (i.e. via liquid biopsy) through detection of genetic signatures of foci that would otherwise be missed with traditional tissue biopsy.

Despite promising initial results, widespread clinical adoption of cfDNA as a biomarker has been impeded by several challenges [4]. One of the most important limitations, especially in the context of variant detection, is the scarcity of circulating tumor DNA (ctDNA) molecules derived from a tumor from typical blood draw volumes, an issue compounded by the weak signal-to-noise ratio of ctDNA with respect to the cfDNA derived from healthy tissue (ctDNA often representing much less than 1% of the total cfDNA fraction) [5,6]. Several strategies have been developed to circumvent this issue, including techniques to enrich tumor derived molecules [7], highly sensitive qPCR-or ddPCR-based assays to detect well-characterized (or personalized) mutations [8–10], and deep sequencing of broad regions of the genome. Each approach has limitations; for example, enrichment techniques are limited to only modest (~2-4 fold) enrichment [7,11] while qPCR-based methods require *a priori* or patient-specific variant knowledge and cannot readily be used for *de novo* discovery or across broad patient cohorts. In some cancers, like prostate, this is especially problematic as even the most common driver mutations exist at frequencies too low to be of broad clinical utility [12]. Targeted deep sequencing, on the other hand, can be used for *de novo* discovery and broader patient coverage, but faces issues concerning sensitivity and specificity introduced by weak tumor signal, clonal hematopoiesis (CH) [13], and technical artifacts introduced during library preparation and sequencing. Additionally, efforts to mitigate these issues are diametrically opposed— at fixed cost, one must choose to either sequence broadly at low depth with reduced sensitivity or more narrowly and deeply but with reduced specificity.

To improve upon detection, we propose a solution that leverages the strengths of targeted deep sequencing and minimizes the weaknesses of traditional panel design by generating a targeted panel guided by machine learning. This solution consists of three strategies: 1) generating a sub exome-sized (2.5Mb) targeted sequencing panel, but instead of only including the coding regions of known cancer genes, focusing on small (~350bp, corresponding to dinucleosomal cfDNA) regions of the genome that are either coding or regulatory non-coding and potentially harbor tumor mutations; 2) computationally selecting candidates for inclusion on this panel with a machine learning model built from actual tumor data and optimized to detect functional or regulatory mutations (“orchid”); and 3) using unique molecular identifiers (UMIs) to suppress technical errors induced by library preparation and sequencing.

In this article, we present our targeted sequencing panel design, demonstrate its *in silico* performance through comparison with two other design approaches, and then validate its ability to detect somatically validated multi-foci tumor variants in the cfDNA of prostate cancer patients at the time of prostatectomy.

## Methods

### Patients Cohorts

This study uses data from two main patient cohorts, including public prostate tumor variant data from 550 patients cataloged in the International Cancer Genome Consortium (ICGC) and 23 (5 for our *in silico* analysis and 18 for our variant capture test) patients from the University of California, San Francisco (UCSF). In the ICGC dataset, patient ages ranged between 32 and 81 (mean of 58.7) and had the following stage distributions: T1 (27%), T2 (39%), T3 (15%), T4 (1%), and Unknown (18%). In the UCSF cohort, patient ages ranged between 50 and 73 (mean of 62.9) and had the following stage distributions: T1 (29.6%), T2 (66.7%), and T3 (3.7%). Additional information about patient cohorts is given in **Supplemental File “Donor Information.xlxs”**.

### Training Data

Whole Genome Sequence (WGS) tumor variant data from the 550 ICGC prostate cancer patients (274 with copy number information) was used to populate a mutation database. In total, the database consisted of 1,588,558 single base substitutions, 66,202 insertions ≤ 200 bp, and 90,255 deletions ≤ 200 bp. Of the 1,717,507 mutations, 90.5% had sequencing coverage between 30-80X. These mutations were annotated with 339 features using the orchid software (http://wittelab.ucsf.edu/orchid) (**‘orchid’ panel; Supplemental**) [14]. Among features, annotations included those related to functional impact, non-coding regulatory status, cancer driver-ness scores, and base-level evolutionary conservation among primates.

### Panel Generation

To build our targeted sequencing panel, we first trained a classification and ranking model, a linear support classifier (SVC), using the orchid software as well as data from our mutation database. We also generated two panels from methods widely used in the field in order to benchmark performance: 1) a gene-centric panel consisting of coding regions from the aggregated set of ~530 genes found in four clinically available cancer-specific targeted sequencing gene panels (referred to as “union-existing”; **Supplemental Table 1**), and 2) a “Frequency” panel, consisting of the most frequent mutations in the ICGC prostate cancer dataset.

### *In Silico* Analysis

We first benchmarked orchid panel’s variant capture performance against the two other designs using an *in silico* analysis of Whole Exome Sequenced (WES) tumor foci DNA and matched cfDNA from 5 patients undergoing radical prostatectomy at UCSF. Somatic variant calling was performed for at least 2 different tumor foci with a normal tissue control for each patient. Next, we generated *in silico* capture probes for the orchid panel by expanding the genomic coordinates of panel mutations by ± 175 bp to match the mode size of cfDNA molecules. Tumor and cfDNA variants were intersected with the orchid panel and the two comparison panels described above.

### Patient ctDNA Variant Detection

The cfDNA from a cohort of 18 prostate cancer patients were isolated and prepared as UMI-tagged libraries for sequencing. After the *in silico* validation of the orchid panel, hybrid capture probes were ordered and used to sequenced panel regions at 2,500X. Tumor variants were subsequently called using the Curio Genomics platform (https://curiogenomics.com), which was designed specifically for processing UMI-barcoded data generated though ThruPlex Tag-seq library preparation by grouping reads into amplification families prior to constructing consensus reads for variant calling. See **Supplemental** for more details.

## Results

### Defining and Evaluating Mutation Classes

It is widely accepted that a relatively small number of genetic variants are responsible for the cellular transformations leading to cancer and that these “drivers” often occur early in a tumor’s evolution leading to high clonality among tumor subclones [15–18]. Additionally, the proportion of drivers decreases relative to the number of passengers as a tumor accumulates mutations [17,19–21]. Following this, we hypothesize that if tumors accumulate mutations as they evolve, those with the lowest mutational burden are both enriched for drivers and more likely to harbor variants at high allele frequency among subclones, making these variants the best candidates for detection in cfDNA.

In order to prioritize which of these low burden mutations should be included on our cfDNA screening panel, we built a mutational scoring model using the sequencing data from the ICGC prostate cancer patients, first defining training labels by dividing ICGC prostate cancer patients into equal-sized groups (n=275 each) based on their number of mutations: 1) Low Burden (LB), consisting of mutations from men with a lower mutational burden, and 2) HB (HB), consisting of mutations from men with a greater burden (**Figure 1A**). We next tested the hypothesis that LB labeled mutations were enriched for drivers, evaluating for their presence in 88 known driver genes (as identified by The Cancer Genome Consortium Prostate Cancer Adenocarcinoma project (TCGA-PRAD) and the IntOGen database; accessed 6/18/2019 [22]) and found significant enrichment in the LB class (hypergeometric test; p = 4.11 e-129), but not the HB class (p > 0.99). This was also the case with 97 prostate driver genes defined by Fraser *et. al.* (LB p = 1.13 e-119; HB p > 0.99) [23]. With these two classes defined, computational complexity was reduced through random down sampling of the data to a total of approximately 50,000 unique mutations while preserving the original LB:HB mutation label ratio in the dataset (approximately 1:40).

**Figure 1:**
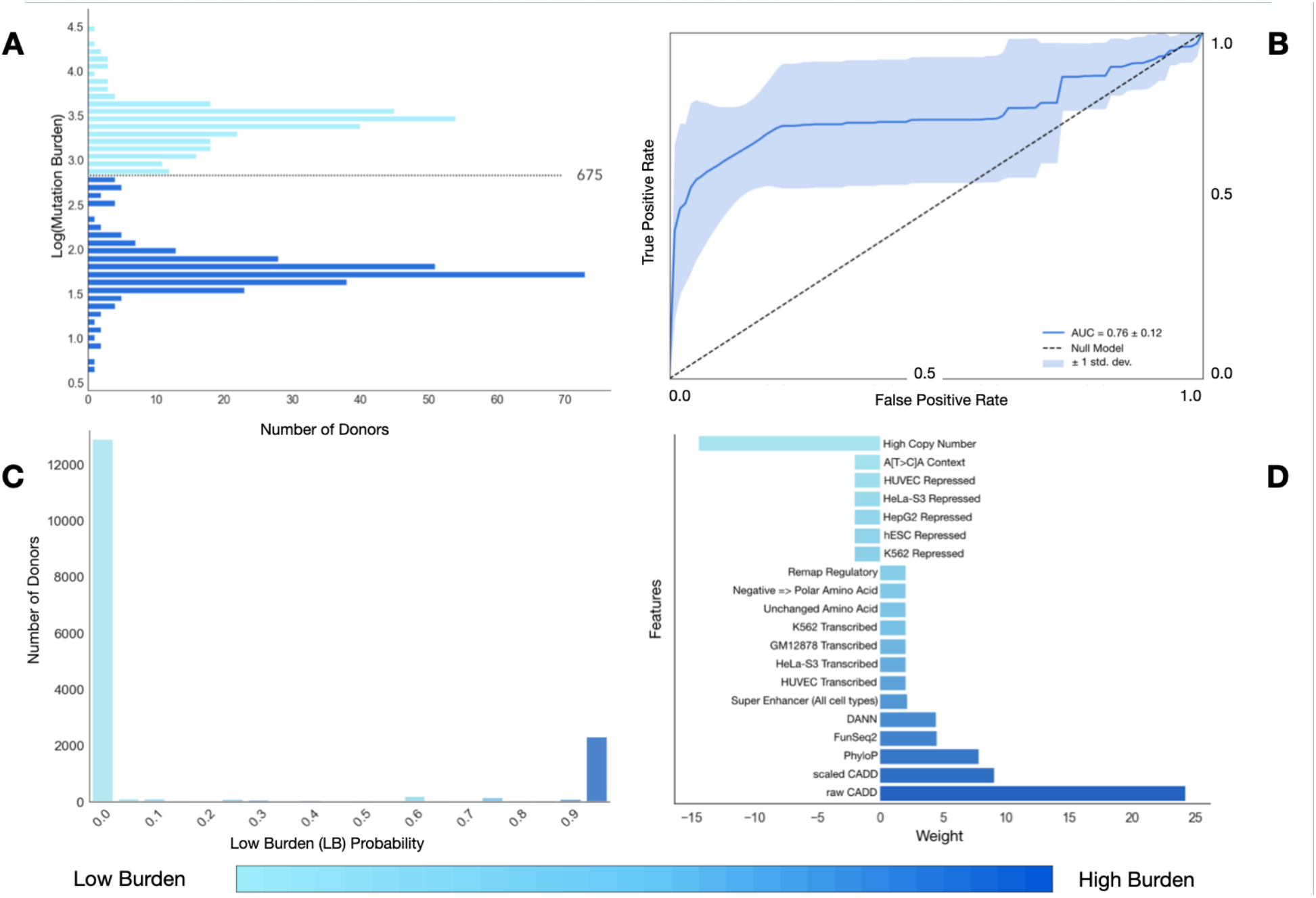
Modeling simple somatic mutations. **A)** We divided ICGC prostate cancer donors into two classes, Low Burden (LB) or High Burden (HB), based on the number of somatic mutations in their tumors and labeled their mutations accordingly. **B)** After modeling with a linear Support Vector Classifier (SVC), we generated a ROC curve of LB classification. Accuracy was 76% +/− 12%. **C)** We visualized classification probabilities for test mutations. The model predicts fewer LB mutations and classifies both LB and HB with high confidence. **D)** We show model feature weights for both classes when features were used as lone predictors. Repressed regions of the genome were more predictive of HB mutations whereas regulatory, transcribed regions of the genome or ‘deleterious’ mutations were more predictive of LB mutations.

### Initial Modeling and Performance

In an effort to guard against overfitting, we used orchid’s feature selection method—which removes features that each account for an average drop in accuracy < 0.1% from the full-featured model—to reduce the number of features from >300 to 20, and performed 10-fold cross validation with a linear SVC, generating a “LB” predictive model. A ROC curve of model performance in the test sets is shown in **Figure 1B**, indicating a 0.76 (± 0.12) classification accuracy. When classification probabilities across all test cases were plotted, we observed a higher likelihood of HB mutation classification, which was expected from the intentional unbalanced LB:HB class ratio used for training (**Figure 1C**). To better understand importance of classification features, we next used each feature singularly in a series new LB/HB classification models, visualizing feature weights and directionality. From this we observed that repressed regions of the genome were predictive of HB mutations, and conversely, regulatory/transcribed regions of the genome—and features indicating strong evolutionary conservation at the base level—were predictive of LB mutations (**Figure 1D**; **Supplemental Figure 1**). This was also expected under our assumption that LB mutations were more likely to be drivers. We elaborate on feature importance in **Supplemental Results**.

### Mutation Ranking

After selecting features, we then built a final classification model fully trained on *down-sampled* data (i.e. none withheld for testing) and used it to score LB probability for *all* prostate cancer mutations in the database. Mutation distances from the fit model’s classification hyperplane were then used to rank them. Those with the greatest magnitudes in the LB direction (i.e. the most “driver-like” or “clonal” under our hypothesis) were further considered for inclusion on the targeted sequencing panel.

### Standardizing Mutation Scores

We annotated candidate LB mutations with associated gene information, if available, using SnpEff [24], as well as functional impact information and transcript length from the UCSC genome database. After binning genes according to their length, we visualized the number of mutations per gene (**Figure 2A**). As expected, longer genes had more mutations (Pearson’s correlation = 0.20, p=6.03 e-39), creating a scenario where marginally scored mutations could be selected for panel inclusion by virtue of strong gene-level feature annotations preferred by the model. To address this issue and to increase gene mutational diversity on the panel, we implemented a corrective standardization (**Supplemental Figure 2**) and applied it to the distance scores of mutations (**Figure 2B**). This standardization reduced Pearson’s correlation between gene length and candidate mutations number to 0.05 (p=1.5 e-3). Mutations that were non-coding or without gene annotation were unaffected by this standardization. After applying this correction, the top 7,034 mutations were then selected for the “orchid” panel. In all, this panel represented 0.41% of the total number of original candidate LB mutations.

**Figure 2:**
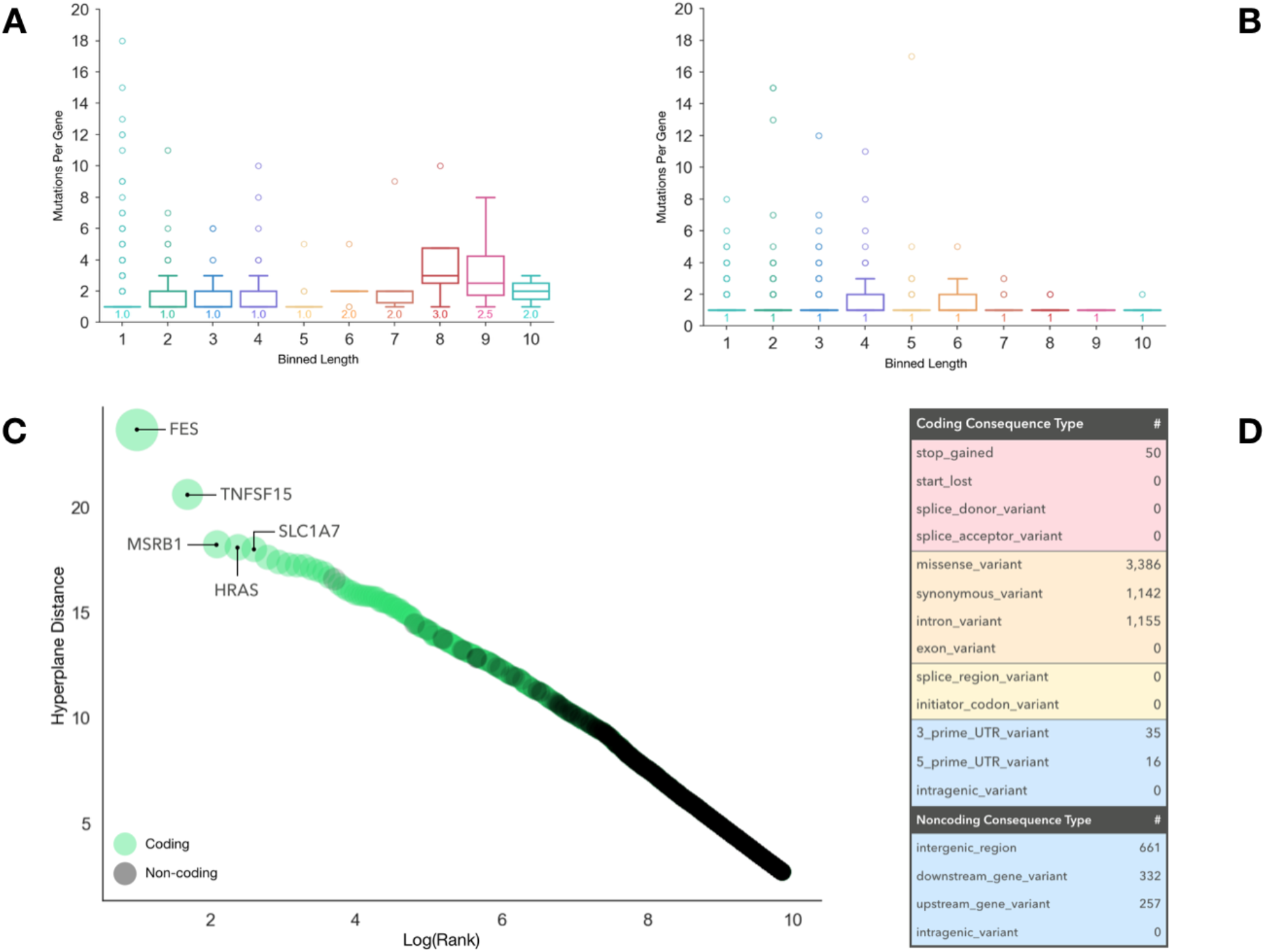
Generating a targeted sequencing library for hybrid capture of LB mutations. We generated a candidate panel consisting of probes targeting the ~7,000 highest ranked LB mutation loci. **A)** We binned genes represented by candidate mutations into 10 groups based on length and show the distribution in number of mutations. Gene length correlated with the number of mutations on the panel (Pearson’s correlation = 0.20, p=6.03e-39). **B)** We employed a distance standardization to mutation hyperplane distances to increase gene diversity on the panel. After standardization the correlation between gene length and number of mutations decreased significantly (Pearson’s correlation = 0.05, p=0.0015). **C)** We plotted the hyperplane distances of retained mutations after standardization against the log mutation rank. Mutations are labeled as coding (green) or non-coding (grey). The top 5 coding mutations with their corresponding genes are labeled. **D)** We show a table of panel mutation consequence types and counts, colored by impact severity (red=high, orange=moderate, yellow=low, blue=modifier).

### Panel Composition

Once our standardized orchid panel was established, we attempted to biologically characterize the mutational composition. Looking at the top 5 coding mutations, for example, we noted that corresponding genes (FES, TNFSF15, MSRB1, HRAS, and SLC1A7) have all been experimentally implicated in cancer as drivers [25,26] (**Figure 2C**). Additionally, we found panel mutations to be significantly enriched for the aforementioned 97 prostate driver mutations (p=2.24 e-13); KEGG-annotated general cancer (p=4.18 e-228) and prostate cancer genes (p=1.19 e-61) (https://www.genome.jp/kegg); and regions associated with regulation of cellular response to growth factors (p=2.68 e-4), MAP kinase activity (p=8.66 e-8), and Integrin signaling (p=1.74 e-13) among others [27–29] (Enrichr; http://amp.pharm.mssm.edu/Enrichr/). Finally, looking at functional impact, we noticed a majority of coding mutations were classified as high or moderate impact and included 50 induced stop gains and 3,386 missense mutations. A table of consequence mutations is shown in **Figure 2D**.

While the most highly ranked mutations were coding, many functional non-coding mutations were also included (~18%) on the panel. For example, we discovered significant enrichment for several general and prostate cancer transcription factor binding sites (**Supplemental Figure 3**), including BRD4 (e=329), CTCF (e=254), FOXA1 (e=188), MYC (e=181), and AR (e=159), as well as a microRNA involved in angiogenesis (mir-126) [27,30] (ReMap; http://tagc.univ-mrs.fr/remap/).

### Panel Performance: *In silico* Analysis

After characterizing the orchid panel, we compared how well it detected somatic variants in relation to two other panels: 1) the union of four existing sequencing panels (Fluxion Biosciences, Foundation Medicine, Guardant Health, and UCSF 500, referred to as the “union-existing” panel; **Supplemental Table 1**); and 2) a frequency-based panel (consisting of the most common mutations; see **Methods**). We assessed this by measuring each panels’ ability to identify somatic tumor-normal variants in multiple tumor foci from 5 prostate cancer patients. Overall, the orchid panel detected more variants than both the frequency panel (p=7.4 e-9) and the union-existing panels (p=3.6 e-10; **Figure 3**), and these differences were statistically significant for all patients except P0024 (only one focus; union-existing [p<0.03], frequency [p<0.02] using a T-test). We also note that, given a fair percentage of the orchid panel (~18%) consists of non-coding regions, tumor variants within these regions could not be assessed through WES data, potentially underestimating the panel’s performance.

**Figure 3:**
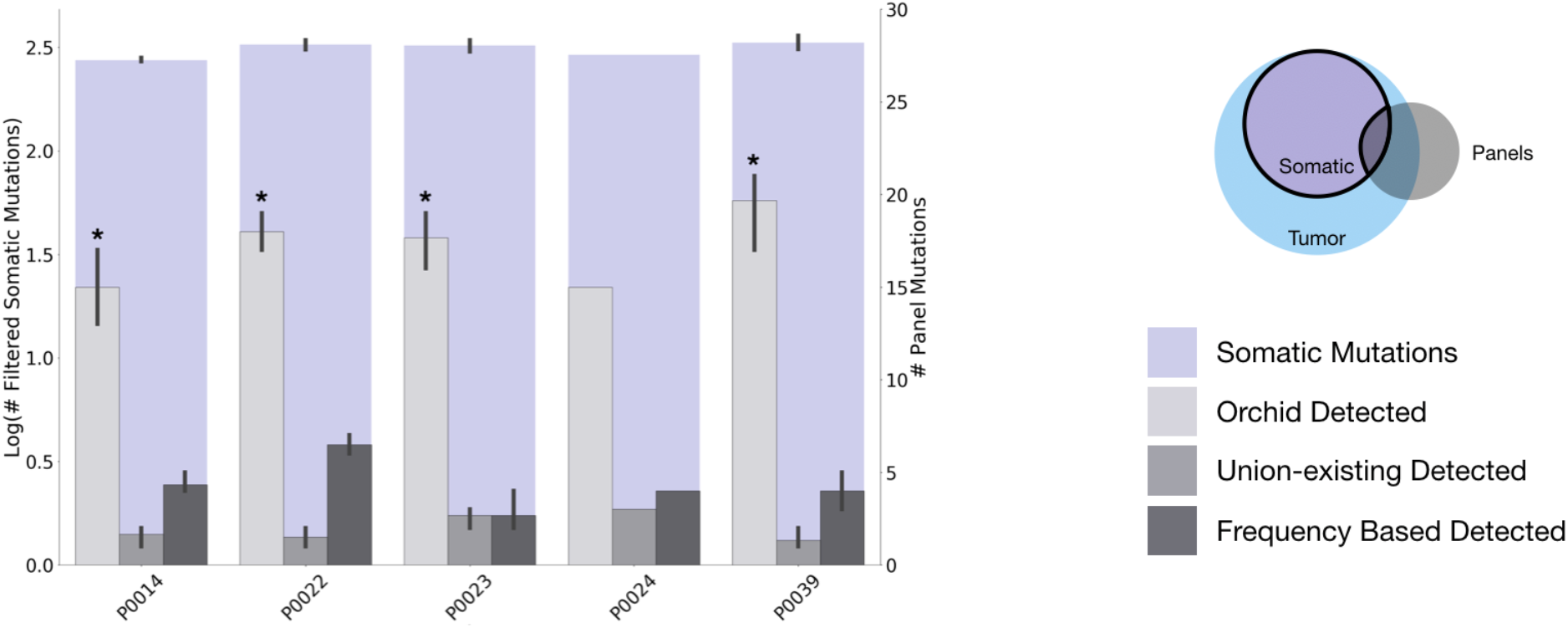
Panel performance using *in silico* capture of cfDNA. Five patients with multiple prostate cancer tumor foci and normal tissue DNA were whole exome sequenced at 200X-fold coverage. Discovered variants were intersected with three *in silico* capture panels: 1) our orchid generated panel, 2) a panel consisting of genes on any of 4 clinically used panels (union-existing), and 3) a panel consisting of all mutations in the ICGC prostate cancer dataset with a frequency > 1 patient. The total log number of somatic mutations from patient foci are shown (purple; left axis) in comparison with those present on each of the three panels (grey-scaled bars; right axis). Orchid detected significantly more mutations in all patients except P0024 (only one focus; union-existing [p<0.03], frequency [p<0.02] using a T-test).

### Panel Performance: ctDNA Variant Detection

After confirming that our orchid-generated machine learning panel improved upon the union-existing and frequency-based panels in an *in silico* setting for detection of mutations in tumor tissue, we ordered hybrid capture probes for regions encompassing the orchid panel mutations (a genomic footprint totaling ~2.5Mb). We then sequenced 18 patients with multiple prostate tumor foci and normal tissues at 2,500X with our panel. Matched cfDNA was also collected for these patients at time of radical prostatectomy and targeted-sequenced at a depth of 2,500X with our panel. We next assessed the panel’s performance in detecting somatic tumor-normal ctDNA variants within the collected cfDNA of these patients. After removing variants not passing quality control filters (see **Supplemental Methods**), we found that variants were detected in all 18 patients, ranging between 15 (S038) and 448 (S076) in number with a median of 122.5. We additionally filtered variants by requiring they be detected in multiple foci of a tumor. In this case, variants were detected in 15 of the 18 patients, ranging between 4 (S025 and S078) and 289 (S076) in number with a median of 26 (**Figure 4A**). The allele frequency of detected variants across patients ranged between 0.24% (S058, S067, and S078; close to the theoretical detection threshold at 2,500X of 0.8%) and 19.82% (S027; a conservative lower threshold for germline variants), with a median of 3.76% (**Figure 4B**). Allele frequency did not correlate with age, stage, or Gleason score (p>0.05).

**Figure 4:**
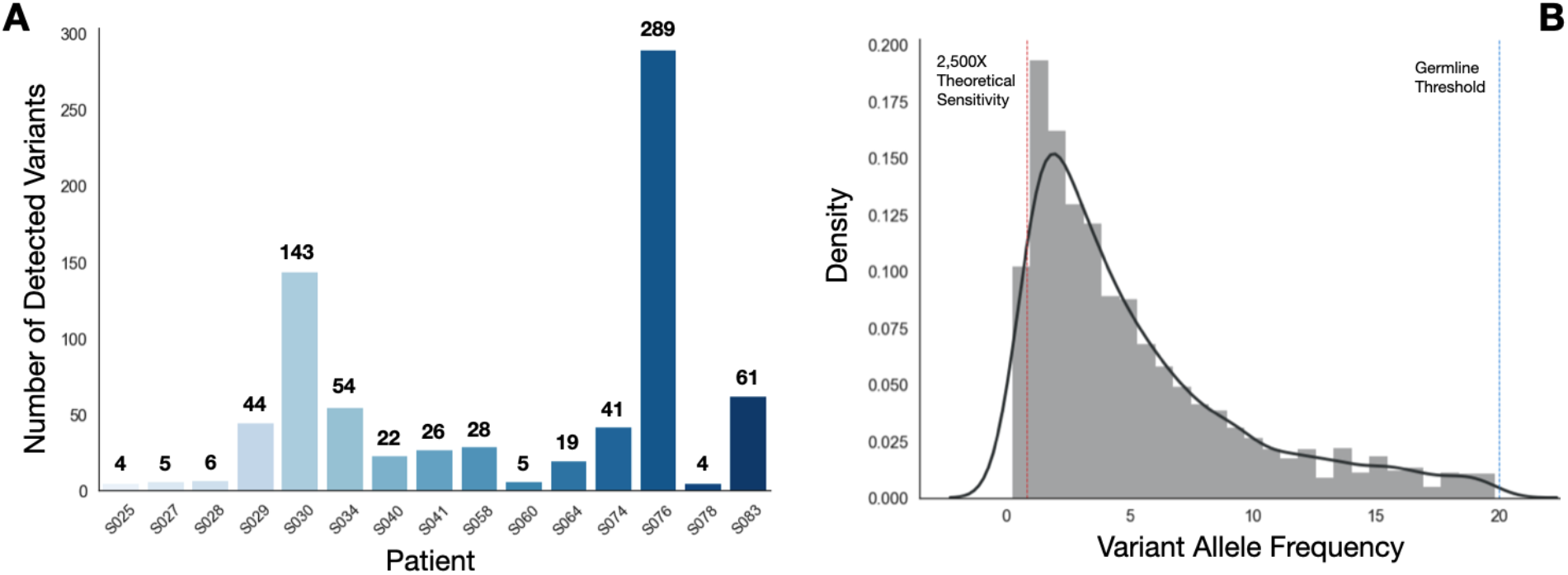
Variant detection and frequency distribution in prostate cancer patients using the orchid generated targeted sequencing panel. Eighteen patients with multiple tumor foci and normal tissue DNA were sequenced at 2,500X-fold coverage after targeted capture using the orchid generated panel. Matched cfDNA was likewise captured and sequenced. **A)** The number of tumor variants detected in the cfDNA of 15 patients is shown. Tumor variants were both somatic and present in multiple tumor foci. Three of the eighteen patients did not have any mutations detected in more than one focus. **B)** The allele frequency distribution of all cfDNA detected tumor variants in A (germline threshold shown at 20%; theoretical sensitivity at 0.8%).

## Discussion

Despite continued progress and marked successes of cfDNA’s application in late-stage disease [2,31,32], ongoing issues prevent wide-spread adoption for early-stage cancer. These issues largely center on tumor heterogeneity and scarcity of tumor derived molecules in circulation. The issues are further compounded by challenges with sample collection and processing, variant artifacts (including CH mutations for non-tumor-matched samples), and bioinformatic analysis. The most straight-forward solutions to mitigate these problems include increasing the volume of blood collected (e.g., 30-100 mL), analyzing ctDNA variants with paired whole blood normal samples, and sequencing at ultra-high depths (e.g. >30,000X). Other solutions include improving molecular techniques, error suppression (e.g. UMIs), and optimizing the composition of gene sequencing panels [11,13,33–37]. Here we expand upon optimizing panels, leveraging machine learning to move past driver-, gene-, or frequency-based panels towards one informed directly by biological datasets. In particular, this is accomplished by modeling low burden mutational signatures developed from tumor/normal sequence and variant annotation data.

While our machine learning approach improved the sequencing panel design, the accuracy of predicting LB *versus* HB mutations was only 0.76. This accuracy can be largely explained by label contamination introduced though incomplete partitioning of driver and clonal variants into the LB class, as presence of these mutations also occurs in the HB class albeit at lower frequency (our hypothesis only assumes *enrichment* in LB). This situation motivated our use of a linear support classifier, which has a higher tolerance of noise (e.g. mislabeled training data) and better feature interpretability relative than other machine learning model types. We found orchid’s feature classification weights to be sensible; for example, associating evolutionary conserved/transcribed regions of the genome with LB tumor mutations, and repressed regions of the genome with HB tumor mutations. Still, despite a fairly high accuracy for noisy data and sensible feature selection, the modeling approach could be improved upon with alternative labeling strategies and/or training data.

There are a number of other qualifications to our machine learning panel design approach that merit consideration. First, establishing a panel’s clinical utility will require much larger sample sizes and greater sequencing depth to further validate variant detection and improve sensitivity in early stage prostate cancer patients. Second, to better elucidate and catalogue CH variants, cfDNA samples should be paired with DNA isolated from whole blood samples and sequenced at equal depth, especially when matched tumor samples are not available. Third, although we compared our panel to two alternative designs *in silico,* future work should compare panels directly using patient samples—ideally with deep whole genome tumor/normal sequence data. Finally, to further assess panel detection as it relates to mutation clonality, follow-up comparison with more sensitive detection strategies (e.g. qPCR), and serial sampling of patient tumors during course of treatment would need to be performed.

## Conclusions

The use of machine learning to optimize targeted sequencing panel composition presents a promising new approach to improve ctDNA variant detection in patients with cancer. In an *in silico* screen, our panel outperformed two alternatives in detecting tumor-derived ctDNA mutations—one generated from a combination of several existing panels, and one based on tumor mutation frequencies. We also demonstrated the targeted panel’s ability to detect tumor variants found in both the cfDNA captured from prostate cancer patients and multiple foci of their tumors.

In summary, we have developed a novel method to rank coding and non-coding tumor mutations for inclusion on a targeted sequencing panel. To our knowledge, this is the first use of machine leaning to generate a capture panel for screening ctDNA of cancer patients. While further research is needed to address the issues of scarce starting material, modeling, and variant discovery, our results provide a useful strategy for broad—yet sensitive—future panel design. Strategies like these are increasingly important for mutation detection in cfDNA isolated from cancer patients with heterogeneous disease, especially at sequencing depths required to reach levels of sensitivity needed for utility in early detection at an affordable cost.

## Supporting information

Donor Information

## Abbreviations

cfDNA: cell-free DNA
ctDNA: circulating tumor DNA
CH: Clonal Hematopoiesis
UMI: Unique Molecular Identifiers
ICGC: International Cancer Genome Consortium
UCSF: University of California, San Francisco
WGS: Whole Genome Sequence
SVC: Support Vector Classifier (linear)
WES: Whole Exome Sequenced
LB: Low Burden
HB: High Burden
TCGA-PRAD: The Cancer Genome Consortium Prostate Cancer Adenocarcinoma

## Declarations

### Ethics approval and consent to participate

All study participants provided informed written consent prior to study enrollment following IRB# 11-05226 approval at the University of California, San Francisco.

### Consent for publication

Not applicable.

### Availability of data and materials

The datasets used and/or analyzed during the current study are available from the corresponding author on reasonable request.

### Authors’ contributions

CC contributed to study design, processed samples, wrote software to generate the screening panel, analyzed and interpreted data, and prepared the manuscript.

EC analyzed and interpreted data.

LL processed samples.

NE interpreted the data.

KL coordinated sample acquisition and processed tumor tissue.

IT coordinated sample acquisition.

JS performed histological examination of tumor tissue and tumor selection for UCSF cohort.

TF coordinated sample acquisition and interpreted data.

PL coordinated sample acquisition.

PP contributed to study design, coordinated sample acquisition, and interpreted data.

PC contributed to study design and acquisition of samples.

JW contributed to study design, interpreted data, and prepared the manuscript.

All authors read and approved the final manuscript.

## Acknowledgements

Not applicable

