## Supplemental for "A machine learning approach to optimizing cell-free DNA sequencing panels: with an application to prostate cancer"

### Supplemental Methods

#### Training Data for the Orchid Panel

#### To build our targeted sequencing panel, we first trained a classification and ranking model using the orchid software [1] and prostate tumor variant data from the International Cancer Genome Consortium (ICGC; <https://icgc.org>; release 23) in conjunction with various biological feature annotation datasets. Briefly, this process consists of annotating each observed mutation with a set of biologically relevant features (e.g. whether the mutation was a non-synonymous amino acid change, present in a DNAse I hypersensitive region, or how evolutionarily conserved, etc.) followed by machine learning on a portion of this data with the goal of classifying a given mutation as originating from a Low Burden (LB) or High Burden (HB) tumor based on this biological feature data. This model was ultimately used to rank variants for inclusion on a custom hybrid capture panel. Using the ICGC Data Portal *Advanced Search* tool, ‘Prostate’ tissue was subsetted from *Primary Site*; ‘WGS’ from *Donor Analysis Type*; and then ‘Simple Somatic Mutation’, ‘Copy Number Somatic Mutation’, and ‘Clinical Data’ data was downloaded and extracted. Data was then imported into a MemSQL database using the *make_database.sh* script from our previously described orchid software. All available feature data (found on the orchid website, <http://wittelab.ucsf.edu/orchid>), was included in database population. Copy number information, when available, was preprocessed and imported using the *parse_cnv.sh* script in the orchid repository. A support vector machine was then trained through orchid as previously described [1].

#### Union of Existing and Frequency Comparison Panels

To benchmark performance of the orchid generated mutation panel in detecting tumor variants, two other panel designs were explored. The first, “Union of Existing”, consisted of coding regions from the aggregated set of 530 genes found from four clinically available cancer-specific targeted sequencing gene panels (**Supplemental Table 1**). This was generated by intersecting gene hg19 coordinates (as queried through Ensembl Biomart at <http://feb2014.archive.ensembl.org/>) with the SeqCap EZ Exome v3 capture panel to include only exons (n=9,470). These regions were then randomly downsampled to match the size of the orchid generated panel (n=7,034). The second comparison panel, “Frequency”, was queried from the ICGC prostate release 23 database. In this case, all mutations present in more than one donor were considered, and ± 175 bp regions centering these mutations were generated to match region sizes used for the orchid panel (n=5,824 regions).

#### Orchid Panel Hybrid Capture Probes

We designed and ordered Custom MyBaits™ hybrid capture probes targeting the final corrected set of mutations, ± 175 bp and tiled 3X through Arbor Biosciences ([https://arborbiosci.com](https://arborbiosci.com/); 16 reaction; catalog 300116). This targeted capture panel was then applied by Arbor Biosciences to library prepped cfDNA samples from Study Sample Patients.

#### Tumor/Normal Sample Extraction and Sequencing

For Initial Cohort samples, multiple tumor tissue foci and normal tissue control samples (seminal vesicle or whole blood if not available) from five stage 1, 2, or 3 prostate cancer patients who underwent radical prostatectomy were collected and processed with the Qiagen DNeasy Blood and Tissue Kit (Catalog # 69504). Samples were sent for whole exome sequencing at 200X. Somatic variant calling was performed using the bcbio GATK workflow (<https://github.com/bcbio/bcbio-nextgen>) to generate raw somatic variants. Tumor and normal variants that did not pass the following filters were initially removed: GATKStandardHaplotypeScore (HaplotypeScore > 13.0), GATKStandardMQ (MQ < 40.0), GATKStandardQD (QD < 2.0), LowQual, GATKStandardReadPosRankSum (ReadPosRankSum < -20.0), GATKStandardMQRankSum (MQRankSum < -12.5), GATKStandardFS (FS > 200.0). Normal variants present in tumor samples were then removed, as well as variants that had associated rsIDs (population level germline mutations). Since the resulting number of filtered tumor variants across all patient/panels was less than 4 and panels were not significantly different from each other (all p-values ≥ 0.6), tumor variants were re-filtered without GATKStandardHaplotypeScore (sensitive to chromosomal abnormalities, a hallmark of cancer), GATKStandardMQ (tumor variants, by nature, have lower mapping quality), and GATKStandardQD (owing to structural changes, quality by depth normalization may be too sensitive) and subsequent germline filtering was performed as above.

For Study samples, multiple tumor foci and normal tissue control samples (seminal vesicle or whole blood if not available) from 18 stage 1,2, or 3 prostate cancer patients who underwent radical prostatectomy were collected and processed with the Qiagen DNeasy Blood and Tissue Kit. Samples were sent for hybrid capture and sequencing at 2,500X. Somatic variant calling was then performed with the SpeedSeq workflow (https://github.com/hall-lab/speedseq) using a minimum of 5% allele fraction, min-repeat-entropy 1, and min-alternate-count 2 for somatic variant calling and default freebayes parameters. Tumor variants were filtered on LOD > 3.5 and tumor/normal read ratio of 2.7 (keeping those flagged with the PASS filter). Only variants that were also present in the cfDNA samples from the same patient were considered.

Tumor tissue samples were obtained using a 2 or 3 mm tissue punch from regions of biopsied prostate that were histologically classified as cancerous with a Gleason scores of 6 or greater by a genitourinary pathologist (JPS). Histologically healthy seminal vesicle (or blood when not available) was also collected to provide a normal sample for background variant analysis.

#### cfDNA Extraction and Sequencing

Between 10 and 20 mL of whole blood was collected from a cohort of prostate cancer patients at time of radical prostatectomy. Blood was first spun at 1,900 g for 10 minutes and collected plasma was re-spun at 16,000 g for 10 minutes to remove any residual cell debris. Samples were then processed using the Qiagen Circulating Nucleic Acid Kit (Catalog # 55114), double eluted with 40 µL of Qiagen Elution Buffer (EB) for 80 µL total and run on the Agilent Bioanalyzer with High Sensitivity DNA chips (Catalog # 5067-4627) to assess concentration and fragment size distribution. Next, for samples meeting a 7.5 ng threshold, the Zymo Clean and Concentrator Kit (Catalog # 4013) was used to concentrate DNA into 10 µL of distilled water and resulting samples were library prepped using the UMI tagging Rubicon ThruPLEX Tag-Seq 48S kit (Catalog # R400585) following kit recommendations for final PCR amplification (7-11 cycles). After AMPure XP bead cleanup (Catalog # A63881) samples were bioanalyzed for quality control and sent to Arbor Biosciences for hybrid capture using the orchid generated panel. Samples used for the orchid variant detection analysis were exome captured with the SeqCap EZ Human Exome Library v3.0 kit, as performed by the Institute for Human Genetics (IHG) as UCSF. In cases where DNA yield was less than that required for sequencing, reamplification for 3 cycles was performed followed by AMPure cleanup and reanalysis via bioanalyzer. Samples were sent to the IHG for sequencing with a target depth of 2,500X (or ~90 Million reads) for panel captured samples and also at 40X for whole exome captured samples.

Sequencing data analysis was performed using Curio Genomics ([www.curiogenomics.com](http://www.curiogenomics.com)) web platform. The following Curio parameters were used: 1) Alignment UMI demultiplexing with a 6 UMI and 11 max stem length, 2) Variant calling with the orchid panel genome feature, 75% family threshold, 2 base hamming distance, and minimum family size of 4 reads, and 3) Filtering with a minimum quality of phred 30, 2 family minimal coverage, and rare allele frequency between 0% and 20%. In some cases, when mentioned, stricter filtering was applied during cfDNA variant analysis. Parameters for this analysis were the same as above except 1) 10 family minimal coverage and 2) 100 total family minimal coverage.

### Supplemental Results

#### Feature Selection and Significance

Sign and magnitude of feature weights define a vector orthogonal to the hyperplane that maximizes the margin between LB and HB classes. Since the dot product of this vector and mutation vectors form the basis of class assignment in a linear SVC, these weights can give insights into their relative importance in the prediction model. Feature weights of the full model (i.e. when all were modeled together) are shown in **Supplemental Figure 1**.

From this information, we observed feature importance patterns that are suggestive of several underlying tumoral mutation pressures. Coding mutations that classified as LB, for example, tended to preserve amino acid identity (‘Unchanged AA’ feature), while also scoring highly on scaled-CADD ‘deleterious mutation’ and PhyloP evolutionary conserved measures. This observation can be explained by a seemingly paradoxical process—tumors with low burden can simultaneously be enriched for a small number of driver mutations (i.e. those altering protein function in favor of growth/survival), while also experiencing strong selective pressures to maintain functionality (via synonymous mutations) of essential growth and survival genes overall (i.e. only tolerating mutations that conserve protein structure) [2].

Interestingly, other mutations with strong evolutionary conservation (or ‘deleteriousness’) scored highly for predicting HB mutations, including FunSeq2, raw CADD, and DANN scores. This discrepancy can be explained by high correlation between feature values—in other words, these features compensate for stronger LB predictors by down weighting overly confident predictions. To test this hypothesis, we remodeled each of these features as single predictors in the full dataset and observed that all ‘deleterious’ features (PhyloP, DANN, CADD raw/scaled, FunSeq2) were predictive of the LB class as shown in **Figure 1D**. The same is true for transcribed/repressed features—when modeled alone, transcribed region features are all predictive of LB mutations, whereas repressed region features mutations are indicative of HB mutations. Future models could make use of better feature selection algorithms that take feature correlation into account.

### Supplemental Figures and Tables

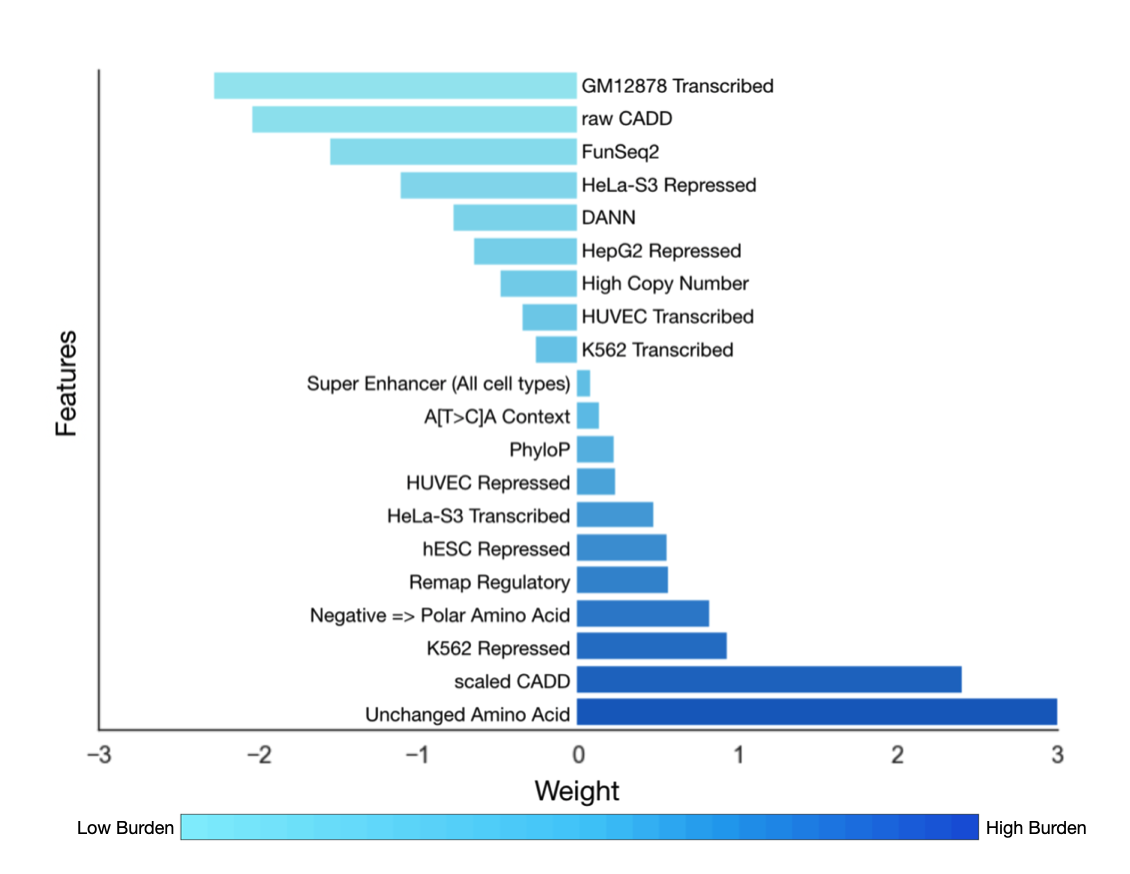

**Supplemental Figure 1: LB/HB Prediction Model with all Features.** We show the SVC model’s feature weights used for the final LB/HB classification and ranking task, indicating relative importance in classification.

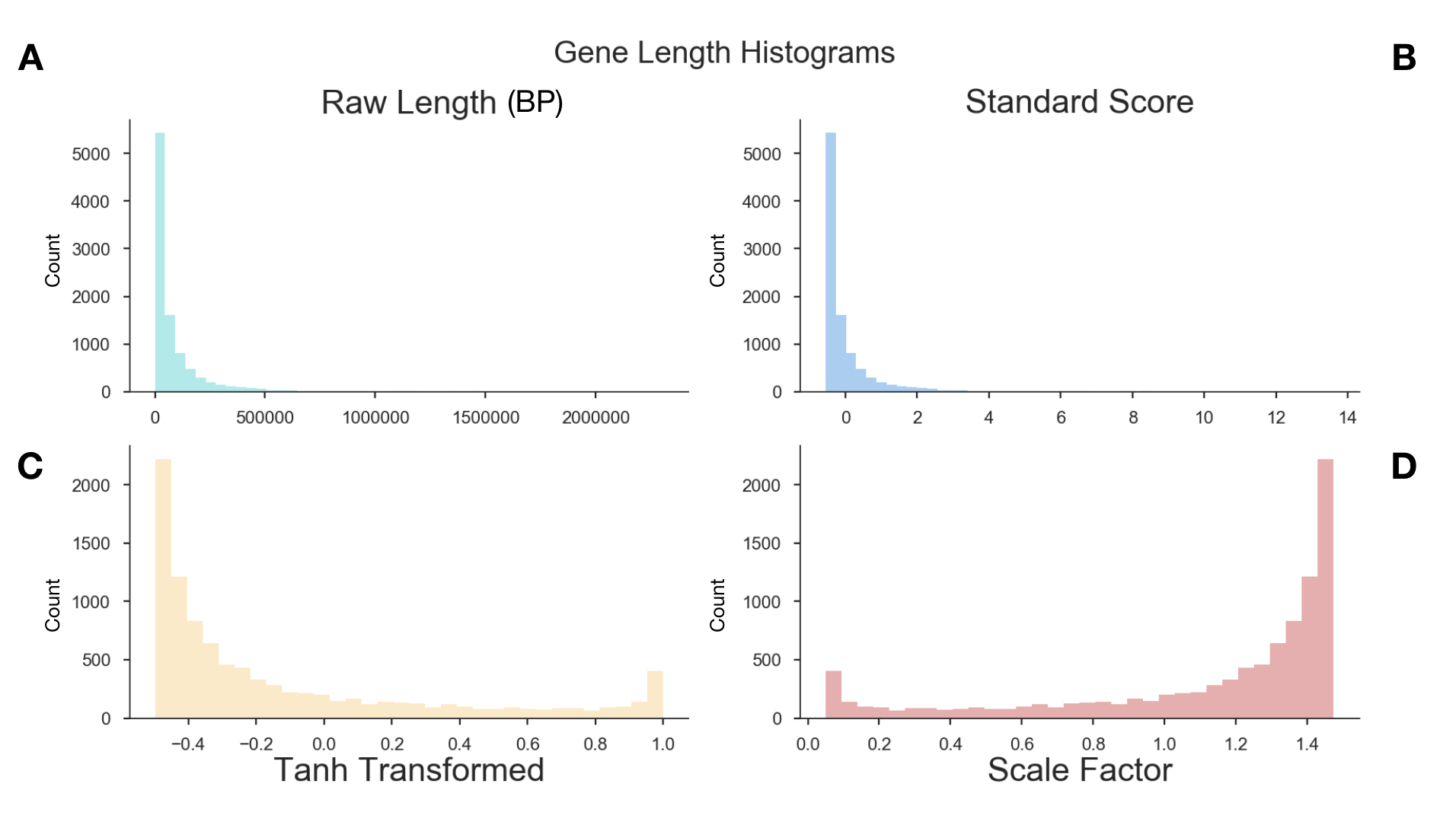

**Supplemental Figure 2: Distribution of gene transcript lengths and scale factors for normalization. A)** A histogram of transcript lengths of genes associated with ranked mutations is shown. **B)** Gene lengths after standardization **C)** Due to the long tail of the distribution of standard scores, a tanh transform was used to compress scores between a -1 to 1 range. **D)** The tanh transformed scores were reversed and multiplied by transcript lengths, effectively down-weighting values of mutations in long genes and up-weighted values of short genes.

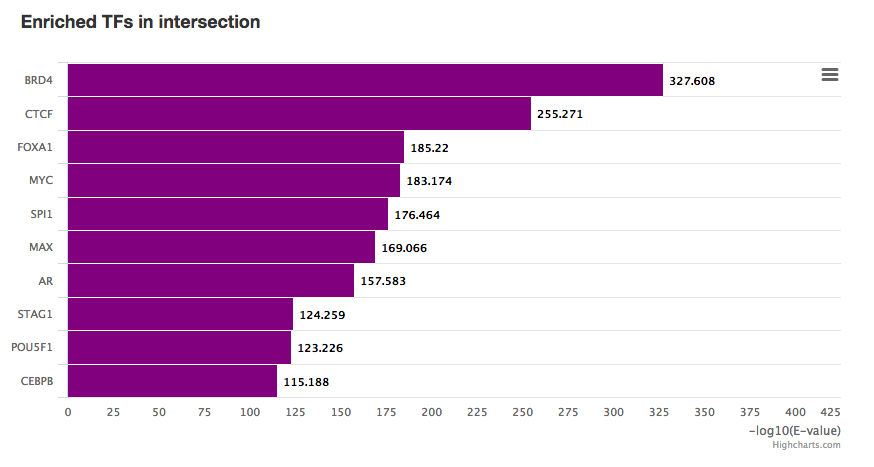

**Supplemental Figure 3: Transcription Factors Enriched on the Orchid Panel.** All non-coding regions on the orchid panel were submitted to the Remap server to predict transcription factor enrichment using 10% overlap with our regions. Shown here are top 10 most enriched transcription factors and their enrichment factors. Most of these are associated with the regulation of cancer, and in some cases, prostate cancer.

**Supplemental Table 1: List of existing panel genes.** Below is the list of genes from 4 combined cfDNA gene panels that were used for the ‘union existing’ panel design in this publication. Genes listed were found from online published material and come from the following panels:

1. UCSF 500: <http://labmed.ucsf.edu/labmanual/db/resource/UCSF_CCGL500_REQUISITION_FORM_2016_withGeneList.pdf>
2. Fluxion Biosciences:

<https://support.fluxionbio.com/hc/en-us/article_attachments/214690287/_634-0042_-_Spotlight_59_Data_Sheet_RevA.pdf>

3. Foundation Medicine: <https://www.foundationmedicineasia.com/dam/assets/pdf/FOne_Current_Gene_List.pdf>
4. Guardant Health:

<http://www.guardant360.com/img/G360MicroSite73GenePanel.jpg>

Note: These panels assess other types of tumor variants beyond simple somatic mutations (including amplifications and gene fusions) and cannot be compared directly to the orchid generated panel for this reason. This list was compiled as a consensus representation of the genes involved in cancer and suggestions a reasonable starting point for defining a targeted sequencing panel.

| ABL1 | C11orf30 | CXCR4 | EZH2 | GNA13 | KDM5A | MLL3 | PAX8 | QKI | SMARCB1 | TSC1 |
| --- | --- | --- | --- | --- | --- | --- | --- | --- | --- | --- |
| ABL2 | C17orf39 | CYLD | FAM123B | GNAQ | KDM5C | MPL | PBRM1 | RAC1 | SMC1A | TSC2 |
| ACVR1 | CALR | DAXX | FAM46C | GNAS | KDM6A | MRE11A | PD-L1 | RAD21 | SMC3 | TSHR |
| ACVR1B | CARD11 | DCC | FANCA | GPC3 | KDR | MSH2 | PD-L2 | RAD50 | SMO | TSHZ2 |
| AJUBA | CBFB | DDIT3 | FANCC | GPR124 | KEAP1 | MSH3 | PDCD1LG2 | RAD51 | SNCAIP | TSHZ3 |
| AKT1 | CBL | DDR2 | FANCD2 | GRIN2A | KEL | MSH6 | PDGFB | RAD51C | SOCS1 | TSLP |
| AKT2 | CBLB | DDX3X | FANCE | GRM3 | KIT | MTOR | PDGFRA | RAD51D | SOS1 | TTYH1 |
| AKT3 | CCND1 | DDX41 | FANCF | GSK3B | KLF4 | MUTYH | PDGFRB | RAF1 | SOS2 | TYK2 |
| ALK | CCND2 | DGKH | FANCG | H3F3A | KLHL6 | MYB | PDK1 | RANBP2 | SOX10 | U2AF1 |
| AMER1 | CCND3 | DICER1 | FANCL | H3F3B | KMT2A | MYBL1 | PHF6 | RARA | SOX2 | USP7 |
| APC | CCNE1 | DIS3 | FAS | HDAC4 | KMT2B | MYC | PHOX2B | RASA1 | SOX9 | VEGFA |
| APOBEC3G | CD274 | DNAJB1 | FAT1 | HDAC9 | KMT2C | MYCL | PIK3C2B | RASA2 | SPEN | VHL |
| AR | CD79A | DNMT3A | FAT3 | HER2 | KMT2D | MYCL1 | PIK3CA | RB1 | SPOP | WHSC1 |
| ARAF | CD79B | DOT1L | FBXW7 | HEY1 | KNSTRN | MYCN | PIK3CB | RBM10 | SPRED1 | WISP3 |
| ARFRP1 | CDC42 | DUSP2 | FGF10 | HGF | KRAS | MYD88 | PIK3CG | REL | SPRY1 | WRN |
| ARHGAP35 | CDC73 | DUSP4 | FGF14 | HIF1A | LEF1 | MYH9 | PIK3R1 | RELA | SPRY2 | WT1 |
| ARID1A | CDH1 | DUSP6 | FGF19 | HIST1H3B | LIFR | MYST3 | PIK3R2 | RET | SPRY4 | XBP1 |
| ARID1B | CDK12 | DYNC1I1 | FGF23 | HMGA2 | LMO1 | NAV3 | PLAG1 | RHEB | SPTA1 | XPO1 |
| ARID2 | CDK4 | EBF1 | FGF3 | HNF1A | LRP1B | NBN | PLCB4 | RHOA | SRC | YAP1 |
| ARID5B | CDK6 | EDNRB | FGF4 | HOXB13 | LYN | NCKAP5 | PLCG2 | RICTOR | SRSF2 | YWHAE |
| ASH2L | CDK8 | EGFR | FGF6 | HRAS | LZTR1 | NCOA2 | PMS1 | RIT1 | SS18 | ZBTB2 |
| ASXL1 | CDKN1A | EGR1 | FGFR1 | HSD3B1 | MAGI2 | NCOA3 | PMS2 | RNF43 | STAG2 | ZBTB20 |
| ASXL2 | CDKN1B | EIF1AX | FGFR2 | HSP90AA1 | MALAT1 | NCOR1 | POLD1 | ROBO1 | STAT3 | ZFHX3 |
| ATF1 | CDKN2A | ELF3 | FGFR3 | HSP90AB1 | MAML2 | NF1 | POLE | ROS1 | STAT4 | ZFHX4 |
| ATM | CDKN2B | EMSY | FGFR4 | HSPA2 | MAP2K1 | NF2 | POLQ | RPL10 | STAT6 | ZMYM3 |
| ATR | CDKN2C | EP300 | FH | HSPA5 | MAP2K2 | NFE2L2 | POT1 | RPTOR | STK11 | ZNF217 |
| ATRX | CEBPA | EPCAM | FLCN | ID3 | MAP2K4 | NFKBIA | POU3F2 | RRAGC | SUFU | ZNF703 |
| AURKA | CHD1 | EPHA2 | FLT1 | IDH1 | MAP3K1 | NFKBIE | PPM1D | RRAS | SYK | ZRSR2 |
| AURKB | CHD2 | EPHA3 | FLT3 | IDH2 | MAP3K2 | NIPBL | PPP2R1A | RRAS2 | SYNE1 |  |
| AXIN1 | CHD4 | EPHA5 | FLT4 | IGF1R | MAP3K5 | NKX2-1 | PPP6C | RSPO2 | TADA1 |  |
| AXIN2 | CHD5 | EPHA7 | FOXA1 | IGF2 | MAP3K7 | NOTCH1 | PRDM1 | RSPO3 | TAF1 |  |
| AXL | CHEK1 | EPHB1 | FOXL2 | IGF2R | MAP3K9 | NOTCH2 | PREX2 | RUNX1 | TBX3 |  |
| BAP1 | CHEK2 | EPOR | FOXO1 | IKBKE | MAPK1 | NOTCH3 | PRKACA | RUNX1T1 | TCEB1 |  |
| BARD1 | CIC | ERBB2 | FOXP1 | IKZF1 | MAPK3 | NPM1 | PRKAG2 | SDHA | TCF7L2 |  |
| BCL2 | CLDN18 | ERBB3 | FRS2 | IKZF2 | MCL1 | NRAS | PRKAR1A | SDHB | TERC |  |
| BCL2A1 | CNOT3 | ERBB4 | FUBP1 | IKZF3 | MDM2 | NSD1 | PRKCA | SDHC | TERT |  |
| BCL2L1 | COL1A1 | ERCC1 | FUS | IL2RB | MDM4 | NT5C2 | PRKCH | SDHD | TET2 |  |
| BCL2L12 | COL2A1 | ERCC2 | FYN | IL7R | MED12 | NTRK1 | PRKCI | SETBP1 | TFE3 |  |
| BCL2L2 | CRCT1 | ERG | GAB2 | INHBA | MEF2B | NTRK2 | PRKDC | SETD2 | TFEB |  |
| BCL6 | CREB1 | ERK1 | GABRA6 | INPP4B | MEK1 | NTRK3 | PRSS8 | SF3B1 | TGFBR2 |  |
| BCOR | CREBBP | ERK2 | GATA1 | IPMK | MEK2 | NUP93 | PTCH1 | SH2B3 | TLR4 |  |
| BCORL1 | CRKL | ERRFI1 | GATA2 | IRF2 | MEN1 | NUTM1 | PTCH2 | SHH | TMPRSS2 |  |
| BLM | CRLF2 | ERRFl1 | GATA3 | IRF4 | MET | OR5L1 | PTEN | SIN3A | TNFAIP3 |  |
| BRAF | CSF1R | ESPL1 | GATA4 | IRS2 | MGA | PAK1 | PTK2B | SLIT2 | TNFRSF14 |  |
| BRCA1 | CSF3R | ESR1 | GATA6 | JAK1 | MGMT | PAK3 | PTPN1 | SLITRK6 | TOP1 |  |
| BRCA2 | CTCF | ESR2 | GID4 | JAK2 | MITF | PALB2 | PTPN11 | SMAD2 | TOP2A |  |
| BRD4 | CTNNA1 | ETS1 | GLI1 | JAK3 | MLH1 | PARK2 | PTPRB | SMAD3 | TP53 |  |
| BRIP1 | CTNNB1 | ETV6 | GLI2 | JAZF1 | MLH3 | PAX3 | PTPRD | SMAD4 | TRAF3 |  |
| BTG1 | CUL3 | EWSR1 | GLl1 | JUN | MLL | PAX5 | PTPRK | SMARCA2 | TRAF7 |  |
| BTK | CUX1 | EZH1 | GNA11 | KAT6A | MLL2 | PAX7 | PTPRT | SMARCA4 | TRIM28 |  |
